# The Representation of Decision Variables in Orbitofrontal Cortex is Longitudinally Stable

**DOI:** 10.1101/2024.02.16.580715

**Authors:** Manning Zhang, Alessandro Livi, Mary Carter, Heide Schoknecht, Andreas Burkhalter, Timothy E. Holy, Camillo Padoa-Schioppa

**Author notes:** Equal contribution (Manning Zhang, Alessandro Livi).

## Abstract

The computation and comparison of subjective values underlying economic choices rely on the orbitofrontal cortex (OFC). In this area, distinct groups of neurons encode the value of individual options, the binary choice outcome, and the chosen value. These variables capture both the input and the output of the choice process, suggesting that the cell groups found in OFC constitute the building blocks of a decision circuit. Here we show that this neural circuit is longitudinally stable. Using two-photon calcium imaging, we recorded from mice choosing between different juice flavors. Recordings of individual cells continued for up to 20 weeks. For each cell and each pair of sessions, we compared the activity profiles using cosine similarity, and we assessed whether the cell encoded the same variable in both sessions. These analyses revealed a high degree of stability and a modest representational drift. A quantitative estimate indicated this drift would not randomize the circuit within the animal’s lifetime.

## Introduction

Economic choices entail computing the values of different options; a decision is then made by comparing values. Work in primates and rodents has shown that these mental processes rely on the orbitofrontal cortex (OFC). Lesion or inactivation of this area severely disrupts choices ^1–4^. Furthermore, experiments using electrical stimulation in monkeys showed that offer values represented in OFC are causal to choices ^5^, and that neuronal activity in OFC directly contributes to value comparison ^6^. Neurophysiology studies examined neuronal activity in monkeys and rodents choosing between different juice types. Neurons in OFC were found to encode different decision variables including the value of individual offers, the binary choice outcome, and the chosen value ^2, 3, 7, 8^. These variables capture both the input (offer value) and the output (choice outcome, chosen value) of the decision process. Moreover, the population dynamics in OFC are consistent with the formation of a decision ^9, 10^. Taken together, these lines of evidence suggest that different groups of neurons identified in OFC constitute the building blocks of a neural circuit in which economic decisions are formed. The organization and the mechanisms governing this circuit are poorly understood. In this perspective, a fundamental question is whether this neural circuit is stable over extended periods of time; alternatively, the representation of decision variables might undergo substantial reorganization ^11^. Importantly, evidence for longitudinal stability would set the stage to study the structure of the circuit, the anatomical identity of different cell groups, and their connectivity.

Longitudinal studies of other neural circuits have reported a spectrum of results. In the parietal cortex of mice engaged in a virtual navigation task, variables encoded by individual cells underwent major reorganization over weeks ^12^. In principle, when variability is confined to a space orthogonal to the coding dimensions, representational drift does not affect the population readout. Interestingly, the drift measured in parietal cortex was constrained by, but not strictly confined to the null space ^13^. Representational drift was also found in visual ^14^ and olfactory ^15^ areas. In particular, longitudinal recordings in the primary visual cortex found relatively stable responses to artificial stimuli (gratings), but substantial drift in the representation of natural scenes ^16–18^. Place fields in the mouse hippocampus were found to be very dynamic (i.e., unstable) ^19^. However, the neuronal propensity to spike was preserved over time ^20^, and changes in spatial tuning were associated primarily with increased experience in the arena, as opposed to the mere passage of time ^21^. In the premotor area HVC of songbirds (zebra finch), the number and timing of spikes of individual neurons in relation to courtship songs remained remarkably stable over weeks ^22^. With respect to OFC, previous work in monkeys found stable representation of decision variables across sessions within a day ^23^. However, it remains unclear whether this representation is also stable over longer periods of time. Importantly, behavioral choice patterns – i.e., the relative values of different goods and the choice accuracy – can vary significantly from day to day ^2, 7^. One hypothesis is that this behavioral variability reflects instability in the decision circuit. Alternatively, choice variability could result from a stable decision circuit and simply reflect variability in the input to that circuit, or changes in the motivational state of the animal.

To shed light on these issues, we used two-photon (2P) calcium (Ca^2+^) imaging in combination with Gradient-Index (GRIN) lenses to record from the OFC of mice performing an economic choice task ^2^. Recordings in each animal continued for many weeks, and we could reliably record from the same cells over many sessions. Neurons were analyzed separately in each session, and the results were compared across sessions. Neuronal activity profiles and the decision variables encoded by individual neurons remained remarkably stable over many weeks. We observed some neuronal changes associated with the degree of experience in the task. In addition, we found a limited representational drift associated with the passage of time. This phenomenon was statistically significant but quantitatively modest. Indeed, the time necessary for a complete reorganization of the neuronal circuit was estimated at >3.5 years – longer than the animal’s lifetime. Thus our results suggest that day-to-day variability in choices is not due to instability in the decision circuit. The presence of a stable circuit within OFC provides new opportunities to examine the mechanisms underlying economic choices.

## Results

### Economic Choices Under the Microscope

Mice were accustomed to head fixation under the 2P objective, and then trained to make binary choices. The choice task closely resembled that used for neurophysiology studies in mice and monkeys ^2, 7^. In each session, the mouse chose between two types of juice labeled A and B (A preferred) and offered in variable quantities. Offers were represented by olfactory stimuli. The odor identity (octanal or octanol) represented the juice type, and the odor concentration represented the juice quantity. For each juice type, we used 4 or 5 quantity/concentration levels. The two odors were presented on the left and on the right, and the animal indicated its choice by licking one of two spouts placed near its mouth (**Fig.1A**). Offered quantities varied from trial to trial pseudo-randomly, and the spatial configuration of the two offers was counterbalanced across trials in each session. **Fig.1B** illustrates the trial structure (see **Methods**).

**Figure 1.**
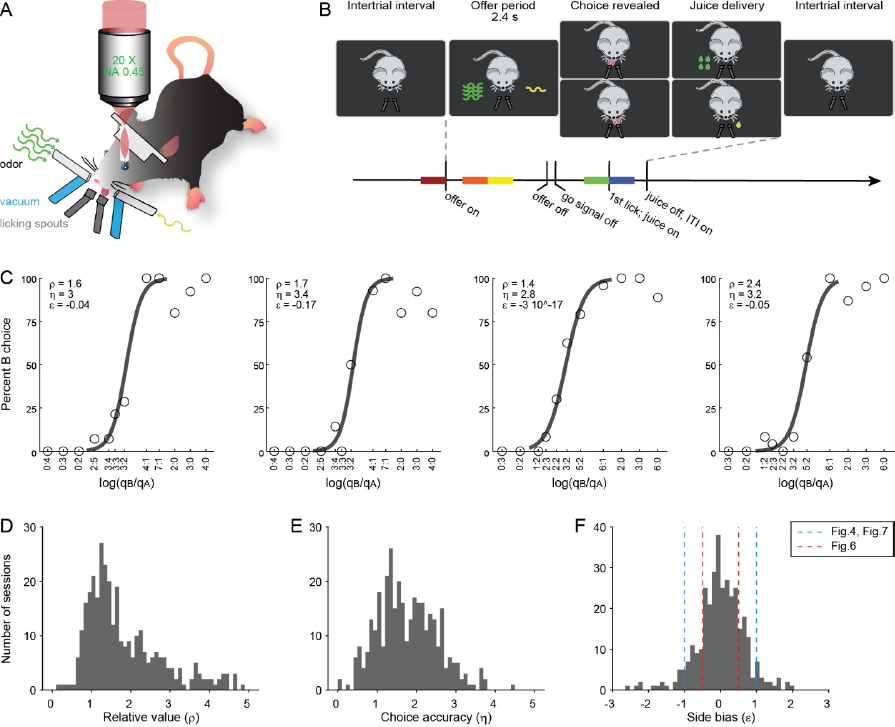
Choice task and behavioral performance. **A.** Experimental apparatus. The animal was head-fixed under the objective. Two odors, each representing one of two offered juices, were presented simultaneously from left and right. The mouse indicated its choice by licking one of two licking spouts. The spatial (left/right) configuration of the offers varied pseudo-randomly from trial to trial. **B.** Trial structure. Following an inter-trial interval (ITI; 2.5-5 s), odors were delivered from the two odor ports. The offer period lasted 2.4 s and ended with an acoustic tone (0.2 s; go signal). The animal had to indicate its choice within 5 s. Immediately after the first lick (following the go signal), the corresponding juice was delivered. During the subsequent ITI, a vacuum system removed from the field all remaining odors. For the analysis of neuronal data, we defined 5 time windows, highlighted here in different colors. **C.** Example sessions, choice patterns. In each plot, the x-axis represents different offered quantities (*q_A_* / *q_B_*). Each data point represents one offer type. The y-axis indicates the percentage of trials in which the animal chose juice B. Sigmoid curves were obtained from logistic regressions (**Eq.2**). Relative value (*ρ*), choice accuracy (*η*), and side bias (*ε*) are indicated for each session. **D-F.** Population histograms for relative value (*ρ*), choice accuracy (*η*), and side bias (*ε*). In panel **F**, dashed lines highlight the inclusion criteria used in the analyses of neuronal data, namely |*ε*| ≤ 1 used for the cosine similarity analysis (Fig.4) and odds ratio analysis (Fig.7) and |*ε*| ≤ 0.5 used for the variable selection analysis (Fig.6) (see **Methods**).

Mice choices reliably presented a trade-off between juice type and juice quantity, as illustrated in **Fig.1C** for four example sessions. For each session, we analyzed choices using logistic regressions (**Methods**, **Eq.1**). This analysis provided measures for the relative value (*ρ*), the choice accuracy (*ƞ*), and the side bias (*ε*). Indicating with *q_A_* and *q_B_* the juice quantities offered in each trial, *ρ* was the quantity ratio *q_B_*/*q_A_* that made the animal indifferent between the two offers; *ƞ* was proportional to the sigmoid steepness and inversely related to choice variability; *ε* was a bias favoring one side over the other (specifically, *ε*<0 and *ε*>0 indicated a bias favoring the left option and right option, respectively). Our entire data set included 343 sessions from 13 animals. **Fig.1D-F** illustrates the distribution for *ρ*, *η*, and *ε* across all sessions. Of note, the side bias was generally modest (|*ε*| < *ρ*) and its distribution across sessions was roughly centered on zero.

We examined whether and how choices changed as a function of the animals’ experience in the task. For each mouse, we counted days starting the first day in which it chose between different juice types. For each behavioral session, we performed a logistic analysis (**Eq.1**), and we derived measures for parameters *ρ*, *η*, and *ε*. For each mouse and for each parameter, we studied how values varied longitudinally by fitting them with an exponential function (see **Methods**, **Eq.3**). **Fig.2A-C** illustrates the results obtained for one example mouse. For this animal, each of the three behavioral parameters showed a significant trend: over sessions, the relative value (*ρ*) decreased, the choice accuracy (*η*) increased, and the side bias (|*ε*|) decreased. We found significant longitudinal effects for 6 of the 12 mice in which we could conduct this analysis, and 4 other mice showed similar trends that did not reach significance level (**Supplemental Table 1**).

**Figure 2.**
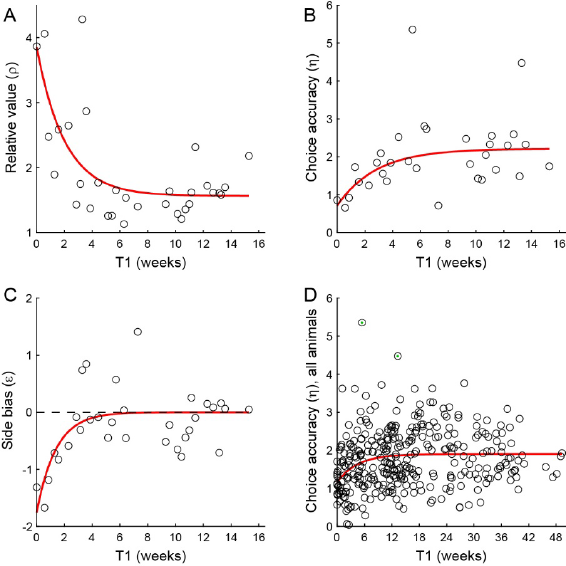
Longitudinal analysis of choices. **A-C.** Results for example mouse (M0032). Relative value (*ρ*, panel A), choice accuracy (*η*, panel B), and side bias (*ε*, panel C) plotted as a function of time. T1 is the time (in weeks) since the first session in which the mouse chose between two juices. Each dot represents one session. For each panel, the red line was obtained from an exponential fit (**Methods**, **Eq.3**). In panel C, the dotted line indicates *ε* = 0. **D.** Longitudinal analysis of choice accuracy. Here, we pooled 343 sessions from 13 mice (entire data set). Each dot represents one session and the red line was obtained from an exponential fit. (Two green dots indicate outliers excluded from the exponential fit.) Referring to **Eq.3**, the fitted parameters (95% confidence interval) were *a* = -0.75 (-1.03, -0.46), *b* = 0.03 (0.005, 0.06) and *c* = 1.90 (1.78, 2.02). Thus the behavioral performance, quantified by the choice accuracy *η*, improved significantly as mice became more expert in the task. The time constant of this trend was τ = 1/*b* = 31 days.

Importantly, the relative value *ρ* and the choice bias *ε* ultimately capture aspects of the animal’s subjective preferences, and longitudinal trends of these parameters do not necessarily indicate a learning process or improved performance. In contrast, in a normative sense, the choice accuracy *η* should be as large as possible (i.e., sigmoids should be as steep as possible) ^24^. Thus for a population analysis we focused on the choice accuracy *η*. We pooled sessions from all animals and we fitted the whole data set with a single exponential function (**Eq.3**). As illustrated in **Fig.2D**, the longitudinal trend was statistically significant. In other words, choice patterns became steeper (i.e., performance improved) over days of experience in the task. The time constant derived from the exponential fit was τ = 31 days.

### Longitudinal 2P Imaging of OFC During Choice Behavior

We conducted 2P Ca^2+^ imaging while mice performed the choice task (see **Methods**). All recordings focused on the left hemisphere. Mice expressing GCaMP6f under human synapsin (hSYN) promoter were implanted with a 1 mm diameter GRIN lens (magnification factor = 1.2) targeting OFC (**Fig.3A**). In each animal, we recorded from multiple fields of view (FOVs). Images from each session were segmented using the CaImAn package ^25^, and the results were checked manually (**Fig. 3BC**). In any FOV, we recorded from 12-225 neurons (median = 77 neurons). Across sessions, we recorded from 2-12 FOVs in each mouse. Thus, by visiting each FOV only once, we could record from 140-850 individual neurons in each mouse.

**Figure 3.**
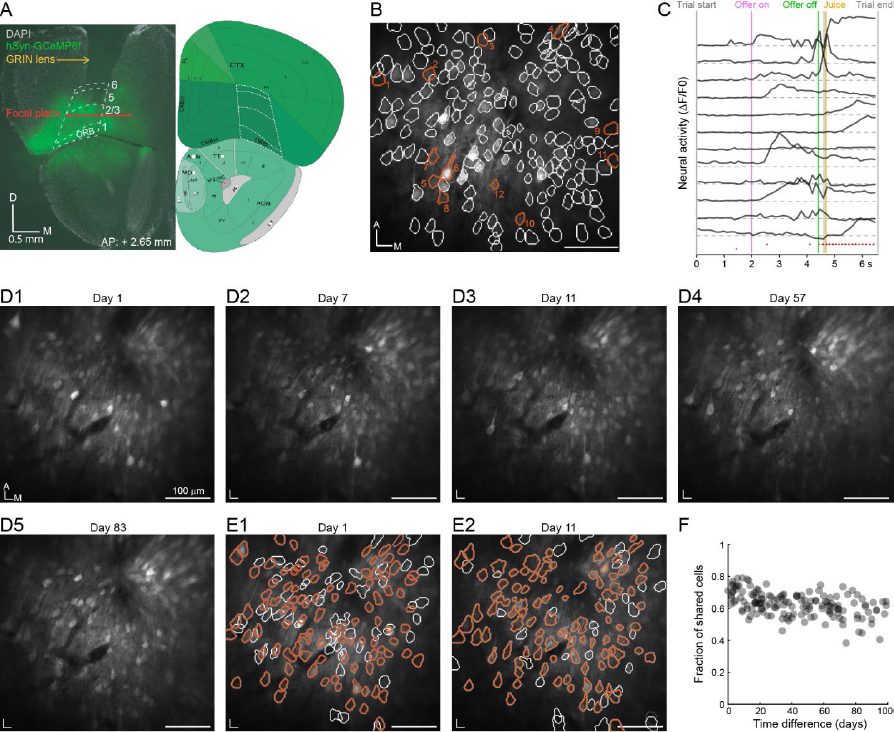
Two-photon imaging of OFC, longitudinal stability. **A**. GCaMP6f viral expression, lens location and focal plane of the lens. Left panel: Coronal slice (AP: +2.65 mm) obtained from a mouse two months after a surgery in which we injected AAV-Syn-GCaMP6f in OFC and implanted a 1 mm diameter GRIN lens. Cell nuclei were stained by DAPI (white), and cells expressing GCaMP6f are contained in the green injection site. The location of the GRIN lens is clearly visible by the lesion in the tissue dorsal to the OFC. The borders of OFC are highlighted (white dotted line) by matching them with a coronal section obtained from Allen Brain Atlas (2011; right panel). The focal plane (red dotted line) was calculated based on the GRIN lens working distance. **B.** Example field of view. Imaging segmentation performed using the CaImAn package ^25^ revealed the presence of N = 169 cells. White contours show neuronal locations. Red contours show 12 example cells illustrated in panel C. 100 μm scale bar and anterior (A), medial (M) directions are shown in the plot. **C**. ΔF/F traces of 12 example neurons (orange contoured cells in panel **B**). Neurons 1 to 12 are from top to bottom. Activity of the 12 cells is shown in one example trial. Vertical lines indicate different behavioral events. Magenta and green shaded areas indicate odor delivery period and reward delivery period, respectively. Red and blue dots show left and right licks, respectively. **D1-D5.** Example FOV recorded in five sessions over the course of 83 days. The day of the first recording was labeled as ‘day 0’ and each image indicates the corresponding recording day. The images remained remarkably stable throughout the recordings. Each image indicates the anterior (A) and medial (M) directions and the 100 μm scale bar. **E1-E2**. Images for day 1 and day 11 obtained after cell segmentation and cell matching. Orange contours indicate matching cells shared between the two FOVs (N = 98 cells identified in both sessions). White contours indicate unshared cells (N = 63 cells for day 1; N = 33 cells for day 11). **F.** Population analysis. We recorded 18 sessions from one single FOV over 100 days. The figure illustrates the fraction of shared cells (number of shared cells/ mean of cell numbers between FOVs) for every pair of sessions (y-axis) as a function of the time difference between the two sessions (x-axis).

Data for the present study were collected from 13 mice and 77 FOVs. For the longitudinal analysis, we recorded from each FOV 2-16 times over a maximum span of 12-41 weeks. For each pair of sessions in which we visited the same FOV, we manually aligned the two sets of images using as landmarks blood vessels and neurons with distinctive features. **Fig.3D** illustrates one example FOV visited 5 times over the course of 83 days. Images across days were remarkably similar. To match individual neurons across pairs of sessions, we used a Bayesian procedure ^26^ that estimated the probability of two cells being the same based on the spatial footprint and the distances from other cells (see **Methods**). Given a pair of sessions, we typically found that 60-80% cells identified in any session could be matched with cells identified in the other session. **Fig.3E** illustrates the results of this analysis for two sessions 11 days apart. Notably, the majority of cells in this FOV could be matched across sessions. Considering all session pairs, the fraction of cells that could be matched across sessions decreased as the number of days between the two sessions increased, but remained >50% even when recordings were 100 days apart (**Fig.3F**).

### Neuronal Activity Profiles Are Longitudinally Stable

After cell segmentation, we normalized the fluorescent trace for each cell and obtained the ΔF/F signal (see **Methods**). All the analyses of neural activity were based on this signal. We first aimed to assess the degree of stability of neuronal activity profiles over time. Referring to the task, a trial type was defined by two quantities (*q_A_*, *q_B_*), their spatial configuration, and a choice. For each session, we examined the ΔF/F trace recorded over the course of each trial. Single-trial traces were obtained joining two time windows aligned with the offer onset (–600 ms, 1600 ms) and with the first lick (–600 ms, 600 ms). We then averaged traces across trials for each trial type. Finally, we concatenated the resulting mean traces across trial types and obtained the activity profile. We repeated this operation for each cell and each pair of sessions, including only trial types present in both sessions. **Fig.4ABC** illustrates the activity profiles recorded for 3 example cells in two sessions at least 7 days apart. Notably, activity profiles differ substantially from cell to cell, but appear relatively stable for each cell. We quantified the likeness between pairs of activity profiles using cosine similarity (CS; see **Methods**, **Eq.4**). In this analysis, activity profiles were vectors where each dimension corresponded to a time bin. CS was the cosine of the angle between two such vectors. In principle, CS could vary between –1 and 1, and CS = 1 if the two traces were identical. For the three example cells illustrated in **Fig.4**, we obtained CS = 0.96, CS = 0.94, and CS = 0.90, respectively.

**Figure 4.**
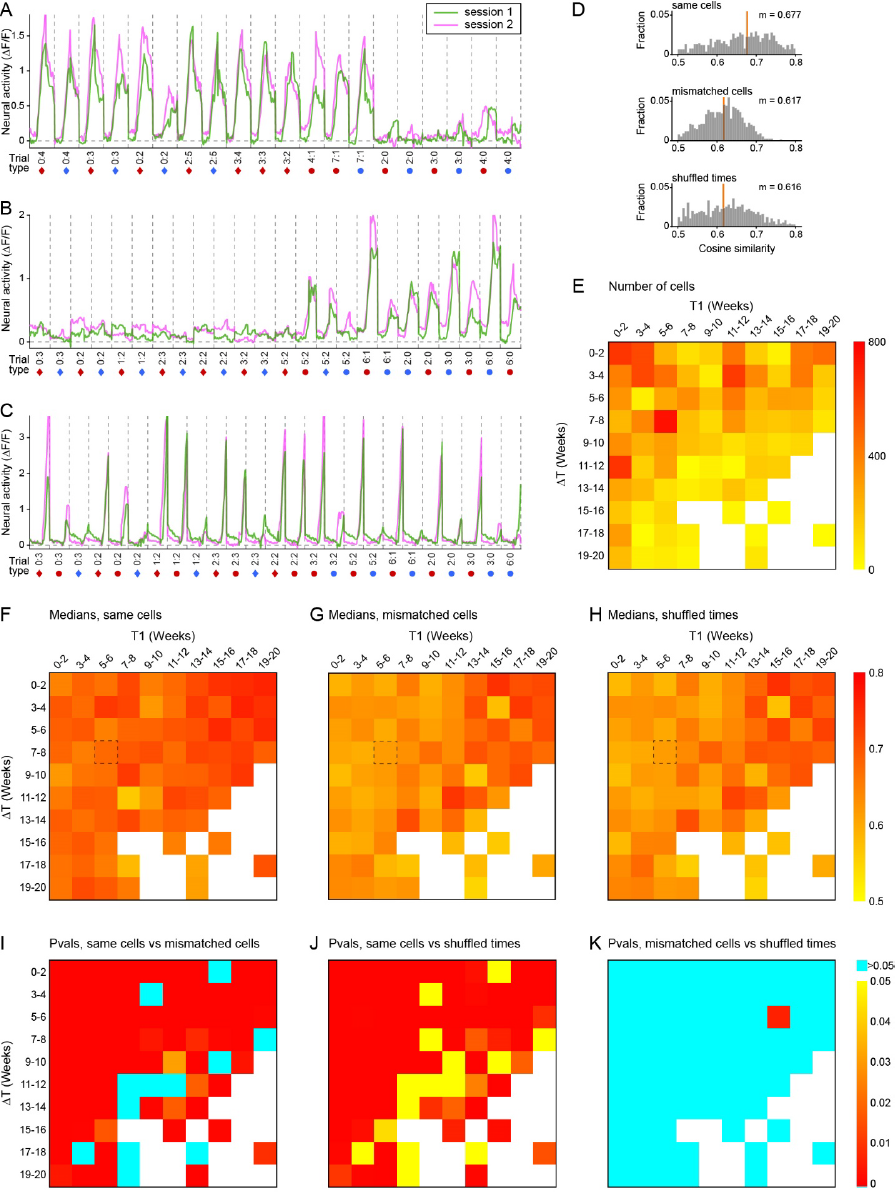
Cosine similarity analysis of activity profiles. **A.** Example cell. We examined the activity of the cell in two sessions (day 31 and day 75). Single-trial traces were obtained by joining two time windows aligned with the offer onset and the time of first lick. In each session, we averaged traces over trials for each trial type, and we concatenated the mean traces obtained for different trial types. The resulting signal is referred to as the activity profile. We repeated this operation for both sessions, and we computed the cosine similarity (CS) between the two activity profiles (see **Methods**). Activity profiles were computed using only trial types present in both sessions. In the figure, activity profiles recorded in session 1 and session 2 are shown in green and pink, respectively. Gray dashed lines separate different trial types. Details about the trial types are indicated below the x-axis: numbers indicate offered quantities (*q_B_*:*q_A_*); diamonds and circles indicate that the animal chose juice A and juice B, respectively; red and blue indicate that the animal chose left and right, respectively. For this neuron, CS = 0.96. **B.** Example cell recorded on days 122 and 129. For this neuron, CS = 0.94. **C.** Example cell recorded on days 1 and 63. For this neuron, CS = 0.90. **D.** Distribution of CS for the three treatments (same cells, mismatched cells, shuffled times). Here T1 = weeks 5-6 and ΔT = weeks 7-8. For each distribution, the orange vertical line indicates the median. We measured median(CS_same_) = 0.677, median(CS_mismatched_) = 0.617, and median(CS_shuffled_) = 0.616. The median measured for the CS_same_ distribution is significantly higher than that measured for the CS_mismatched_ distribution (p = 2.26 10^-38^, Kruskal-Wallis test) and that measured for the CS_shuffled_ distribution (p = 1.37 10^-29^, Kruskal-Wallis test). **E.** Number of cells matched across sessions as a function of T1 and ΔT. Session pairs were divided in 2 weeks x 2 weeks bins (see main text). Here, the number of cells available in each bin are illustrated with a heat map (see color legend). Each entry in the table indicates the number of cells recorded with the corresponding T1 (x-axis) and ΔT (y-axis). Bins with ≤10 cells were removed from this figure. **FGH.** Measures of median(CS) as a function of T1 and ΔT. The three panels refer to three treatments, namely same cells (**F**), mismatched cells (**G**), and shuffled times (**H**). In each panel, median(CS) is illustrated with a heat map. The black dashed rectangle highlights the bin illustrated in panel D. The color bar on the right is for all three panels **F**, **G**, **H**. **IJK.** Statistical comparison of different treatments. Panel **I** illustrates the p-values obtained comparing the distributions of CSs measured for “same cells” and “mismatched cells” treatments. Panel **J** illustrates the p-values obtained comparing “same cells” and “shuffled times” treatments. Panel **K** illustrates the p-values obtained comparing “mismatched cells” and “shuffled times” treatments. All p-values were obtained from a Kruskal-Wallis test. The color bar on the right is for all the three panels (**I**, **J**, **K**).

To assess how CS varied across the neuronal population and as a function of time, we adopted the following terminology and procedures. Starting from the first day of recording, for any two sessions, T1 and T2 indicated the two recording days (with T1≤T2; see **Methods**). We also defined the time separation ΔT = T2 – T1. We examined how CS varied as a function of both T1 (i.e., experience in the task) and ΔT (i.e., time passage). First, we considered the plane defined by T1 and ΔT and we partitioned it in two-dimensional time bins of 2-weeks x 2-weeks. Both T1 and ΔT ranged between 0 and 20 weeks (100 time bins total). Second, for each time bin in this plane, we considered neurons that had been recorded in two sessions, at the corresponding times T1 and ΔT in the plane. Because of the large number of FOVs included in this study and the multitude of recording sessions obtained for each FOV, our data set included a considerable number of cells for most bins (**Fig.4E**). Third, taking advantage of this rich data set, we examined the distribution of CS obtained in each time bin.

**Fig.4D** illustrates this analysis for one representative time bin (T1 = weeks 5-6; ΔT = weeks 7-8; N = 747 cells). Notably, the distribution of CSs was broad and almost entirely >0.5 (median CS = 0.677). Since our measures of ΔF/F were nearly always >0, profile vectors were essentially confined to a single orthant of a high-dimensional space, which implies that CS was necessarily large. To gauge whether our measures of CS were higher than one might expect by chance, we examined the distribution of CSs obtained for two control treatments. First, for any FOV recorded in two sessions, we arbitrarily mismatched the cell pairings by randomly coupling neurons from session 1 to neurons from session 2 (treatment: mismatched cells). We then proceeded with the computation of CSs and generated the resulting distribution. As expected, CSs were generally >0.5. However, the median of this distribution (median CS = 0.617) was significantly lower than that measured when neurons were matched (p = 2.26 10^-38^; Kruskal-Wallis test). Second for any cell recorded in two sessions, we arbitrarily shuffled the activity profile by randomly permuting the components of the profile vector (treatment: shuffled times). Again, CSs obtained for this treatment were generally >0.5. However, the median of this distribution (median CS = 0.616) was also significantly lower than that measured for the original activity profiles (p = 1.37 10^-29^; Kruskal-Wallis test). We concluded that the activity profiles measured for the same neuron in these two sessions were significantly more similar than expected by chance.

We repeated these analyses for each time bin. Thus for every pair of T1 and ΔT, we obtained three measures of median(CS) corresponding to the three treatments (same cells, mismatched cells, and shuffled times). Heat maps in **Fig.4FGH** illustrate the results of these analyses. Several points are noteworthy.

First, the median(CS) measured for same cells was higher than that computed for either of the two control treatments. This was true for every value of T1 and ΔT, and the difference was statistically significant for most values of T1 and ΔT (**Fig.4IJ**; Kruskal-Wallis test). In other words, the activity profiles recorded for individual neurons in different sessions were significantly more similar to each other than expected by chance.

Second, a direct comparison of the two control treatments indicated that, for most time bins, the median(CS) obtained for mismatched cells and that obtained for shuffled times were statistically indistinguishable from each other (**Fig.4K**). Importantly, the time shuffling procedure is non-biological and only preserves the fact that ΔF/F is generally >0. Thus one might expect that the median(CS) obtained for the time shuffled control would be substantially lower than that obtained for mismatched neurons. Conversely, the fact that the CS distributions obtained for the two control treatments were statistically indistinguishable indicates a high degree of variability in the activity profiles of different cells (aside from the fact that ΔF/F was generally >0).

Third, visual inspection of **Fig.4F** suggested that median(CS) increased as a function of T1 and decreased as a function of ΔT. To assess the validity of this observation, we defined larger time bins of 6-weeks x 6-weeks (**Fig.5A**). We refer to these larger bins as Bins. We then compared the CS distributions measured for different Bins. (1) To ascertain whether activity profiles varied as a function of T1, we compared CSs measured in Bin 1 (T1 = weeks 0-6; ΔT = weeks 0-6) and in Bin 2 (T1 = weeks 15-20; ΔT = weeks 0-6). Confirming our impression, median (CS) was significantly higher in Bin 2 than in Bin 1 (p = 1.55 10^-115^, Kruskal-Wallis test). In other words, the activity profiles of individual neurons became more reproducible as the animals became more experienced in the task. (2) To ascertain whether activity profiles varied as a function of ΔT, we compared CSs measured in Bin 1 (T1 = weeks 0-6; ΔT = weeks 0-6) and in Bin 3 (T1 = weeks 0-6; ΔT = weeks 15-20). Indeed, median (CS) was significantly lower in Bin 3 than Bin 1 (p = 0.05, Kruskal-Wallis test). In other words, the activity profiles of individual neurons became more dissimilar as a function of the time distance between sessions (representational drift).

**Figure 5.**
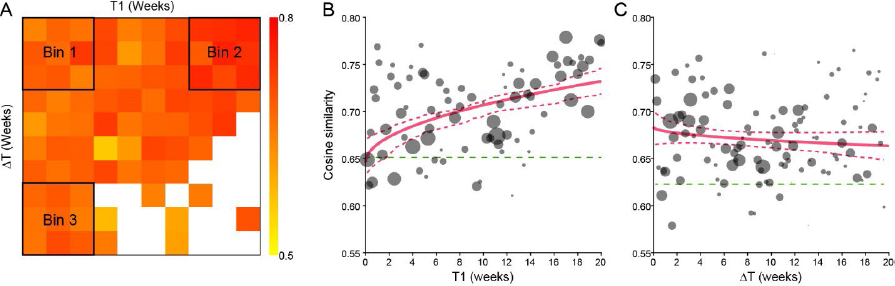
Cosine similarity, longitudinal trends. **A.** Median(CS) plotted as a function of T1 and ΔT (same cells). This panel is as in Fig.4F, and we highlighted the three 6 week x 6 week Bins used for longitudinal comparisons. Median(CS) was significantly higher in Bin 2 than in Bin 1 (p = 1.55 10^-115^, Kruskal-Wallis test); and significantly lower in Bin 3 than Bin 1 (p = 0.05, Kruskal-Wallis test). **B.** Cosine similarity increases with task experience (T1). This analysis was restricted to pairs of sessions with ΔT ≤ 6 weeks, so that we had a comparable number of cells for different values of T1 (see Fig.4E). For each day, we computed the media(CS). In this plot, the radius of each circle is proportional to the number of cells. The green dashed line indicates the median(CS) obtained by mismatching cells. The red line was obtained from the fit CS(T1) = *a_0_* + *a_1_* T1^1/2^. Red dashed lines indicate the 95% confidence interval. **C.** Cosine similarity decreases with time passage (ΔT). Same conventions as in panel B. This analysis was restricted to pairs of sessions with T1 ≤ 6 weeks. We fitted the data with a drift model CS(ΔT) = *a_0_* + *a_1_* ΔT^1/2^, and we obtained *a_0_* = 0.6835 and *a_1_* = -1.687 10^-3^ day^-1/2^. We also computed *b* = median(CS_mismatched cells_) for the same population, and we obtained *b* = 0.622821 (green dashed line). Thus we estimated the time necessary for full reorganization, namely ΔT_FR_ = 1280 days.

To further characterize these longitudinal trends, we examined how CS varied as a function of T1 (experience in the task) and ΔT (time passage) with a 1 day temporal resolution (**Fig.5BC**). Motivated by an analogy between representational drift and a diffusion process in which displacement increases as the square-root of time, we quantified the representational drift with the regression CS(ΔT) = *a_0_* + *a_1_* ΔT^1/2^, from which we obtained *a_0_* = 0.6835 and *a_1_* = -1.687 10^-3^ day^-1/2^. We also computed *b* = median(CS) for the “mismatched cells” treatment. In this context, *b* can be regarded as the chance level for CS – i.e., the expected value for CS if the population underwent full reorganization. We obtained *b* = 0.622821. On these bases, we estimated the time necessary for full reorganization of the neural circuit given the drift rate, namely ΔT_FR_, as follows:

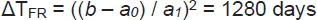

In other words, the time necessary for a full reorganization was >3.5 years – longer than the animal’s lifetime. In conclusion, the representational drift observed in this neuronal population was statistically significant but quantitatively modest.

Control analyses based on alternate measures of similarity and/or restricted to task-related cells yielded similar results (**Supplemental Figure 1**; **Supplemental Figure 2**).

### Neuronal Encoding of Decision Variables

We next examined the variables encoded by individual neurons in OFC. This analysis was similar to that previously conducted for spiking activity in monkeys and mice ^2, 7^. We defined five time windows aligned with different behavioral events: pre-offer (0.6 s before the offer onset), post-offer (0.4–1 s after the offer onset), late delay (1–1.6 s after the offer onset), pre-juice (0.6 s before juice delivery onset), post-juice (0.6 s after juice delivery onset) (**Fig.1B**). Again, a trial type was defined by two offered quantities, their spatial configuration, and a choice. For each cell, each time window, and each trial type, we averaged ΔF/F over trials. A neuronal response was defined as the activity of one neuron in one time window as a function of the trial type. Each FOV was analyzed based on a single session (i.e., each neuron contributed only once to this analysis). Thus our data set included 6620 neurons from 13 animals and 73 FOVs (see **Methods**).

The analysis proceeded in steps. First, we examined the activity of each cell in each time window with an ANOVA (factor: trial type). We set a threshold of p<0.01. Neurons that satisfied this criterion in ≥1 time window were identified as task-related. A total of 932 cells were task-related and included in subsequent analyses. Second, we examined how neuronal responses varied as a function of the trial type. Qualitatively, different responses seemed to encode different variables, including the value of individual offers, the spatial configuration of the offers, the chosen value, and the chosen side. **Fig.6A-D** illustrates four representative examples.

**Figure 6.**
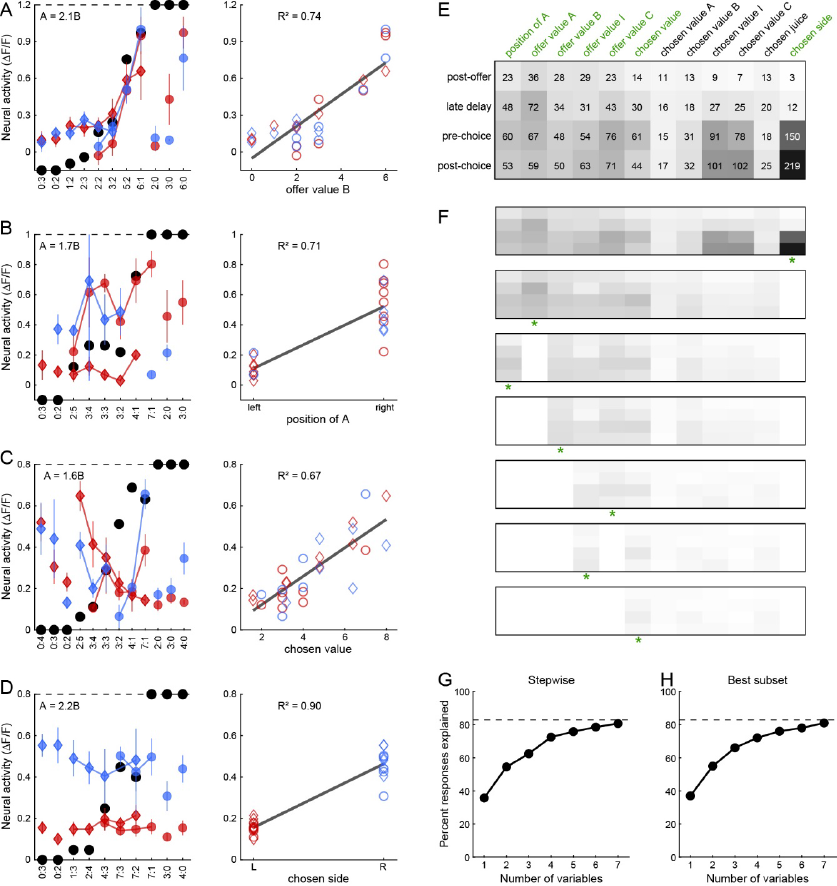
Decision variables encoded in OFC. **A.** Example cell encoding the *offer value B*. In the left panel, offer types are ordered by the quantity ratio *q_B_*/*q_A_* (x-axis). Black dots represent the choice pattern. The relative value (ρ = 2.1), derived from logistic analysis, is indicated in the panel. Color symbols represent the neuronal activity (ΔF/F). Diamonds and circles are for trials in which the animal chose juice A and juice B, respectively. Red and blue are for trials in which the animal chose the offer on the left and right, respectively. In the right panel, the same neuronal response is plotted against the binary variable *offer value B*. The cell activity increased as a function of *q_B_*. The dark gray line was obtained from a linear regression (R^2^ indicated in the panel). B. Example cell encoding the *position of A*. Same conventions as in panel A. The activity of this cell was roughly binary – low when juice A was offered on the left, and high when juice A was offered on the right. **C.** Example cell encoding the *chosen value*. **D.** Example cell encoding the *chosen side*. Neuronal responses illustrated here are from (**A**) late delay, (**B**) late delay, (**C**) post-juice, and (**D**) pre-juice time windows. **E.** Population analysis. Each response passing the ANOVA criterion was regressed against each of the 12 variables defined in **Supplemental Table 2**. If the regression slope was non-zero, the variable was said to explain the response. Any given response could be explained by more than one variable, and the best-fit variable was identified as that providing the highest R^2^. The panel illustrates for each time window (row), the number of responses best explained by each variable (column). Here, numbers and shades of gray indicate cell counts. **F.** Variable selection, stepwise procedure. The top image is the same as in panel **E**. In the first iteration, we identified the variable that explained the highest number of responses (*chosen side*). We removed from the pool all the responses explained by that variable (35.73% of the total). The residual pool is illustrated in the second image. The procedure was then repeated. At each iteration, we identified the variable that explained the largest number of responses in the residual pool (highlighted by a green asterisk). If the marginal explanatory power was ≥2%, we retained the variable and removed from the pool all the responses explained by that variable. The procedure continued until all newly selected variables failed the 2% criterion. **G.** Percent of explained responses, stepwise procedure. The x-axis indicates the number of variables (iterations). On the y-axis, 100% corresponds to the number of responses passing the ANOVA criterion; the dotted line corresponds to the number of responses collectively explained by the 12 variables included in the analysis. **H.** Percent of explained response, best-subset procedure. Same format as in panel **G**. In panel **E**, variables selected by both the stepwise and best-subset procedures are highlighted in green.

For a quantitative analysis, we defined twelve candidate variables possibly encoded by neurons in OFC (see **Methods**, **Supplemental Table 2**). We then regressed each task-related response separately against each candidate variable. If the regression slope differed significantly from zero (p<0.05), the variable was said to explain the response and we noted the corresponding R^2^. If a response was explained by more than one variable, we identified the variable providing the best explanation as that with the highest R^2^. Thus we generated a population table for the number of responses best explained by each variable in each time window (**Fig.6E**).

We aimed to identify a small number of variables that would explain most of the neuronal data set. As for the analysis of spiking data, we adopted two procedures for variable selection, namely stepwise and best-subset ^7,27^. The stepwise procedure identifies at each iteration the variable that explains the largest number of yet unexplained responses, and then removes from the data set every response explained by that variable. In the first 3 iterations, this procedure selected variables *chosen side*, *offer value A*, and *position of A* (**Fig.6F**). We continued the procedure until when any additional variable explained <2% of responses (**Fig.6G**). Selected variables included *offer value A*, *offer value B*, *position of A*, *offer value left*, *offer value right*, *chosen value* and *chosen side*. Importantly, the stepwise procedure is path-dependent and does not guarantee optimality. In contrast, the best-subset procedure identifies for k = 1, 2, 3, … the subset of k variables that collectively explain the maximum number of responses (i.e., yield the maximum explanatory power). For k = 7, the best-subset procedure selected the same variables selected by the stepwise procedure (**Fig.6H**). Notably, the variables identified here match those found in the analysis of spiking data (see **Discussion**).

Finally, we classified each neuron by assigning it to the variable providing the largest total R^2^ across time windows. This sum was restricted to the time windows for which the cell passed the ANOVA criterion. Task-related cells that were not explained by any variable were classified as *untuned*. Subsequent analyses on the longitudinal stability of neuronal functional roles were based on this classification.

### The Encoding of Decision Variables is Longitudinally Stable

We aimed to assess whether individual neurons maintained their functional role – i.e., whether they encoded the same variable over extended periods of time. For each FOV, we selected two sessions based on the behavioral performance and the number of cells (see **Methods**). We labeled them as session 1 (earlier) and session 2 (later). We identified neurons that had been recorded in both sessions and that were task-related in both sessions. We repeated this operation for all FOVs, and we obtained a data set of 649 cells. We then examined the classification obtained for each neuron in each session.

Qualitatively, the responses recorded for any given neuron in the two sessions often appeared very similar (**Supplemental Figure 3**). For a population analysis, we constructed a contingency table where rows and columns represented the variable encoded in session 1 and in session 2, respectively, and each entry was the corresponding number of neurons (**Fig.7A**). In this table, cells maintaining their functional role would populate the main diagonal. Conversely, neurons encoding different variables in the two sessions would be off diagonal. Neurons that encoded one variable in one session and that were *untuned* in the other session would populate the rightmost column or the bottom row.

**Figure 7.**
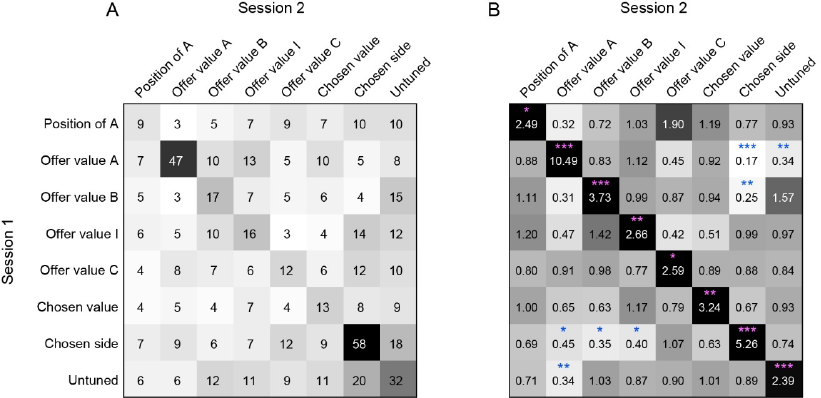
The encoding of decision variables is longitudinally stable. **A.** Contingency table. Each neuron was recorded and classified in two sessions. Task-related cells not explained by any variable were classified as *untuned*. Each entry (*i* , *j*) of this contingency table indicates the number of cells classified as encoding variable *i* in session 1 (rows) and variable *j* in session 2 (columns). **B.** Table of odds ratios (ORs). Numbers represent the OR obtained for each entry in in panel A. The chance level is OR=1, and OR>1 (OR<1) indicates that the cell count was higher (lower) than expected by chance. For each location, we performed Fisher’s exact test to assess whether the departure from chance level was statistically significant. Pink and blue asterisks indicate that the cell count was significantly above and below chance, respectively (* p<0.05, ** p<0.01, *** p<0.001).

Because the seven variables were encoded with different frequencies (different numbers of cells), the contingency table in **Fig.7A** is not immediately interpretable. We thus generated the corresponding table of odds ratios (ORs; **Fig.7B**). In this table, each entry (*i*, *j*) quantifies how the corresponding cell count departs from that expected by chance, given the total number of cells encoding variable *i* in session 1 and variable *j* in session 2 (see **Methods**, **Eq.7**). For each entry, OR = 1 is the chance level and OR > 1 (OR < 1) indicates that the cell count was higher (lower) than expected by chance. Inspection of **Fig.7B** reveals that for all diagonal entries cell counts were significantly >1 (Fisher’s exact test). In other words, the neuronal representation of each variable in OFC was significantly more stable than expected by chance. A complementary aspect of the results emerges upon inspection of the rightmost column and bottom row of **Fig.7B**. A reasonable intuition might be that, over the course of many weeks, some neurons become tuned to the task while other neurons leave the pool. Contrary to this hypothesis, all entries in the rightmost column and bottom row of the OR table were at or significantly below chance level. In other words, cells were typically tuned in both sessions, or not at all. In conclusion, the results illustrated in **Fig.7B** indicated that the encoding of decision variables in OFC was very stable. These findings appear particularly notable if we consider the fact that the variables encoded in OFC are correlated with each other, and that measures of neuronal activity are intrinsically noisy.

**Fig.7** included only two sessions for each FOV. However, our data set was substantially richer, because we recorded from the same FOV 2-16 times over the span of 12-41 weeks (depending on the animal and the FOV). Thus we conducted a longitudinal assessment of encoding stability. For each FOV and for each pair of sessions, we generated a contingency table and an OR table similar to those displayed in **Fig.7**. Encoding consistency was ultimately captured by the diagonal nature of the OR table. To quantify this trait, we used the diagonalization index (DI; **Methods**, **Eq.8**). We examined how DI varied as a function of the time distance (ΔT). Our analyses indicated (1) that for any ΔT, the consistency in classification was significantly higher than that expected by chance and (2) that DI did not significantly decrease as a function of ΔT (**Supplemental Figure 4**).

Importantly, measures of mean(DI) suggested that our classification procedures were noisy. Indeed, our success rate can be estimated at mean(DI)^1/2^ ≈ 55%. A previous study in monkeys ^23^, using a similar approach had provided an estimated success rate of ≈ 80%. The difference might be due to two factors, namely (1) that the mouse OFC represents more variables (making the classification more challenging), and (2) that mice completed fewer trials each day. For any given level of neuronal noise, the indeterminacy in the tuning function decreases with the number of trials.

## Discussion

Neurophysiology studies in primates and rodents suggested that different groups of neurons in OFC constitute the building blocks of a decision circuit. To investigate the structure and thus the mechanisms of this circuit, we developed a protocol for 2P Ca^2+^ imaging in mice performing an economic choice task. Imaging of OFC was performed through GRIN lenses. The protocol provided very stable signals, in the sense that we were able to record from the same FOV and from the same neurons repeatedly for up to 40 weeks. Taking advantage of this preparation, we examined the degree of longitudinal stability in neuronal activity profiles and the representation of decision variables. Several results presented in this study are particularly noteworthy.

First, we explicitly examined longitudinal changes in behavioral choice patterns. An earlier study noted that capuchins (a species of New World monkey) can effectively choose between pairs of foods never encountered together before ^28^, suggesting that economic choice is a natural behavior. Numerous studies in other species^29–31^ and anecdotal observations in macaques support this view. For example, when we train monkeys in a standard juice choice task ^7^, we familiarize animals with the trial structure using only one juice; subsequently, we introduce choices between two different juices. Typically, animals present non-trivial trade-offs (i.e., 1<*ρ*<∞) from the very first session in which two juices are offered against each other, consistent with the notion that economic choice is a natural behavior. Importantly, this notion does not exclude that performance in the choice task might improve with experience, although – to our knowledge – this question has not been addressed previously. Here we conducted a longitudinal analysis to that effect. Ultimately, task performance is quantified by the choice accuracy *η*, which is proportional to the steepness of the behavioral sigmoid ^24, 32^. For given relative value *ρ*, a larger *η* (i.e., a steeper sigmoid) means that there are fewer trials in which the animal chooses the lower value. Our analyses showed that the choice accuracy increased over sessions, and reached a plateau with a time constant estimated at 31 days. In principle, multiple factors might contribute to this behavioral trend. (1) While choosing between two juices, mice must decode the stimuli representing the two offers – i.e., two odors presented in variable concentrations on the two sides of the nose. Although mice have a sophisticated olfactory system, it seems likely that the perceptual component of the task was somewhat challenging for them, and that over the course of sessions animals improved in their ability to decode the olfactory stimuli. (2) Aside from specific components, performance in our task certainly required some degree of mental focus. It is quite possible that, over sessions, mice became increasingly capable of focusing on the choice task for many trials. This point is particularly salient if one considers the fact that in binary choices there is nominally no error – i.e., even when they choose the lower value, animals still receive some juice quantity at the end of the trial.

Supporting the notion of increasing mental focus, the number of trials any animal could perform in any one sitting typically increased with experience. (3) Last but not least, the decision mechanisms might become more efficacious over time. Training improves performance in a variety of natural functions including perception and motor control; it seems reasonable to assume that training also improves performance in decision making. An important question for future research concerns the relative weights of these three factors.

Second, we identified a set of variables encoded by individual neurons in OFC. These include variables associated with individual offers (*offer value A*, *offer value B*, *offer value ipsi*, *offer value contra*), with the spatial configuration (*position of A*), and with the choice outcome (*chosen side*, *chosen value*). This finding strongly resonates with the results of neurophysiology studies in primates ^7^ and mice ^2^. One difference is that the set identified here includes both juice-based and spatial variables, while previous work in mice had found a prevalence of spatial variables ^2^. We will address this apparent discrepancy in detail in a separate report.

Third, longitudinal analyses provided evidence for high stability in the activity profiles of individual cells and in the representation of decision variables. Analyses based on cosine similarity revealed that activity profiles became gradually more reproducible as the animals became more experienced in the task – a phenomenon that parallels behavioral observations. In addition, activity profiles drifted as a function of time passage. However, this drift was quantitatively modest, and the estimated time necessary for full reorganization exceeded the lifetime of the animal. Concurrently, analyses based on the decision variables did not reveal any systematic trend. Considering the fact that only a fraction of OFC neurons appear as task-related in any given session, a reasonable hypothesis was that, over the course of many weeks, some neurons would leave the active pool while other neurons would become tuned to the task. Another reasonable hypothesis held that the functional role of any given cell would evolve over time. Our results argued against both hypotheses. Notwithstanding the fact that classification procedures are somewhat noisy, neurons in OFC were typically tuned consistently across sessions, or not tuned at all. Furthermore, neurons typically encoded the same variables across sessions. These results strongly support the notion of a stable decision circuit. Consequently, they set the stage to examine the connectivity of this circuit and, ultimately, the mechanisms governing the winner-take-all process underlying economic decisions.

It is interesting to consider our findings in the context of a growing literature on representational drift. Previous studies reported a broad spectrum of results. Substantial drift was found in visual ^14–18^, olfactory ^15^, and parietal ^12^ areas. In the hippocampus, place fields were found to drift substantially ^19, 20^, but this phenomenon was related to the experience in the arena as opposed to the passage of time per se ^21^. Finally, in the premotor HVC region of zebra finches, the representation of courtship songs was found to be very stable ^22^. Our results place the representation of economic decision variables in OFC on the stable side of the spectrum. Under the assumption that brain regions controlling high cognitive functions are more flexible and/or less rigidly organized ^33^, our findings might appear somewhat counterintuitive. At the same time, the fact that decision variables in OFC are encoded categorically^34–36^ likely contributes to the stability of this neuronal representation. Indeed, a neuron drifting from one decision variable to another would have to undertake a quantum leap, qualitatively different from what might take place, for example, in primary visual cortex. In a computational perspective, some degree of stability seems necessary for a circuit performing a winner-take-all process. Thus, in our opinion, the most significant aspect of the results is that they lay the ground to investigate the organization of this decision circuit. Future research shall address this fundamental question.

## Methods

All experimental procedures were approved by the Washington University Institutional Animal Care and Use Committee (IACUC).

### Animals and surgical procedures

The study was conducted on N=13 mice (6 males, 7 females) expressing the genetically encoded Ca^2+^ indicator GCaMP6f under a pan-neuronal promoter. N=10 C57BL/6J animals (B6; Jackson Laboratory, stock #000664) were injected the pAAV.Syn.GCaMP6f.WPRE.SV40 (Addgene, stock #100837) virus. N=2 Rasgrf2-2A-dCre animals (Jackson Laboratory, stock #022864) and N=1 PV-Cre animal (Jackson Laboratory, stock #008069) were injected a 1:1 mixture of pAAV.Syn.GCaMP6f.WPRE.SV40 (Addgene, stock #100837) virus and pAAV.CAG.LSL.tdTomato (Addgene, stock #100048) virus. In every mouse strain, we aimed to express GCaMP6f under synapsin promoter. During the imaging experiments, animals were 2.5-7 months old.

In a single surgery, we injected the virus(es), implanted a gradient-index (GRIN) lens, and fixed a head bar to the skull. During surgery, anesthesia was provided by 0.8-1.5% isoflurane in 2 L/min medical air. A heating pad maintained the body temperature at 37°C. Buprenorphine Sustained-Release (SR) (1 mg/kg, subcutaneous (s.c.) injection) was injected for long-term (2 days) pain relief, and dexamethasone (2 mg/kg, intramuscular (i.m.) injection) was injected to limit inflammation. We also injected 2% lidocaine hydrochloride locally at the incision site to reduce pain. Sterilized saline (37 °C, 0.50 ml, intraperitoneal (i.p.) injection) was injected at the end of the surgery to avoid dehydration. We first retracted the skin overlying the left and right hemisphere of the mouse to expose the skull overlying frontal and parietal cortices. We then performed a 2-3 mm diameter craniotomy spanning OFC. **Supplemental Table 3** details the target locations in OFC for each mouse. After removing the skull, we injected the GCaMP6f virus (or a mix of GCaMP6f and LSL-tdTomato viruses) into the brain at two locations (AP: +0.15 mm, ML: -0.15 mm; reference point: target location) and (AP: -0.15 mm, ML: +0.15 mm, reference point: target location) with a Hamilton syringe (Hamilton stock #65458-02). At each location, we injected 100 nl in the tissue at DV depth ranging -0.1 to +0.1 mm with 0.05 mm step size (reference point: target location). The injection speed was 20-40 nl/min and we waited 3 minutes between steps. Because this type of syringe has a bevel at the needle tip, we corrected the injection depth by adding 0.7 mm (= distance between the tip of the needle and the opening’s center). After withdrawing the needle, we slowly inserted a 1 mm diameter GRIN lens (Inscopix ProView lens probe, stock #1050-004605, magnification factor = 1.2, numerical aperture (NA) = 0.5 in air) into the brain and down to OFC (AP: 0 mm, ML: 0 mm, DV: -0.15 to - 0.3 mm; reference point: target location). The 0.15-0.3 mm (see **Supplemental Table 3**) were subtracted to correct for the fact that this GRIN lens has a working distance ranging 100-300 μm. This procedure was performed with a GRIN lens holder (Inscopix) at the speed of 0.6 mm/min. During lens insertion, to give some opportunity for the brain to rebound, we retracted 0.2 mm upon first reaching depths of 1, 1.5, and 2 mm. Once in place, the GRIN lens was secured to the skull using C&B Metabond dental cement (Parkell stock #S380). Finally, we attached a custom head bar to the skull using dental cement. After surgery, mice were given dexamethasone (2 mg/kg, i.m.) for inflammation and carprofen (5 mg/kg, s.c.) for pain, if needed. Animals were given 7-14 days to recover before we started training or imaging sessions.

### Histology

To verify the locations of GRIN lens implantation and GCaMP6f expression, after completing all the in vivo imaging experiments, we performed trans-cardiac perfusion ^37^ to harvest the mouse brains. Mice were anaesthetized deeply with ketamine/xylazine cocktail (10 mg/Kg, i.p. injection) and then perfused with phosphate-buffered saline (PBS) (mixed with heparin with ratio of 100:1) and 4% paraformaldehyde (PFA). We removed the head bar and GRIN lens. Brains were then extracted from the skull and embedded into 15% then 30% sucrose/PBS solution at 4° C for several days, until they sunk into the bottom of the solution. We then covered the brains with O.C.T. compound (Tissue-Tek) and froze them at -80° C. Brains were sliced in 40 μm thick coronal sections using cryostats (Leica Biosystems, CM1950) and cover-slipped with antifade mounting medium with DAPI (Vector Laboratories, VECTASHIELD). Brain slices were imaged through inverted microscopy (Leica DMI6000B) with 2.5X, 5X and 10X objectives (Leica Plan Apo). Slice imaging results were compared to the Allen Mouse Brain Atlas (Coronal Atlas) from AP: +2.60 mm to +2.80 mm. We used the lesion gap caused by the GRIN lens as a landmark to check whether recorded FOVs were within OFC, accounting for the GRIN lens range of working distances.

### Choice task and training protocol

The choice task closely resembled that used for non-human primates ^7^ and was nearly identical to that described in a previous mouse neurophysiology study ^2^. During training or imaging sessions, mice were head-fixed and placed in a customized tube under the microscope. In each session, the animal chose between two different juices labeled A and B (juice A preferred) and offered in variable quantities. In each trial, we delivered two odor stimuli (octanal and octanol) simultaneously from the left and right of the mouse nose (**Fig.1A**). The odor identities represented the juice type, and the odor concentration represented the juice quantity. For juice A, we used 5 different concentrations to represent 5 quantities (1-5 drops). For juice B, we used 4 or 5 different concentrations to represent 4 (2,3,4,7 drops) or 5 quantities (2-6 drops or 1,2,3,5,6 drops), respectively. The association between odors and juices was balanced across mice (in 8 animals, juices A/B were paired with octanal/octanol, respectively; in 5 animals, the association was reversed). The offered quantities were linearly related to the odor concentration (e.g., odor levels 1 to maximum number of ppm linearly represented quantity 1 to maximum number of juice). Offered quantities *q_A_* and *q_B_* (and the corresponding odor concentrations) varied from trial to trial. For any two quantities, the spatial configuration of the offers (left/right) varied pseudo-randomly from trial to trial.

Odors were presented for 2.4 s (offer period). Immediately thereafter, a 0.2 s tone was played (go signal). In each trial, the animal indicated its choice by licking one of two spouts. The animal had to indicate its choice within 5 s. The lick was detected by a battery-operated touch circuit ^38^, and the chosen juice was immediately delivered. Juice delivery typically took 150-1050 ms (∼150 ms/per drop). If the animal did not respond within 5 s, the trial was aborted. Licks preceding the onset of the go signal were ignored and did not result in any juice delivery. In forced choice trials (∼35% of trials), only one odor was presented on one side (one correct response). If the animal licked the wrong spout, a 2-5 s white noise sound was played. The exact length of the white noise sound varied across sessions but was fixed within each session. Between two trials, we activated a vacuum system that removed the odors remaining from the previous trial. The inter-trial interval (ITI) varies in the range 2.5-5 s.

On any given day, mice typically performed the choice task for 30 mins (1 session; 170-270 trials) and received ∼0.7-1.3 ml liquid. Juice pairs were varied across mice and included: (A) 15% sucrose vs. (B) water; (A) apple juice vs. (B) blueberry juice; (A) apple juice vs. (B) water; (A) 15% sucrose vs. (B) cranberry juice; (A) 15% sucrose vs. (B) grape juice; (A) water vs. (B) apple juice; (A) pomegranate juice vs. (B) water; (A) apple juice vs. (B) white grape juice; (A) white grape juice vs. (B) water; (A) grape juice vs. (B) cranberry juice; (A) 15% sucrose vs. (B) cranberry juice; (A) 15% sucrose vs. (B) peppermint tea; (A) elderberry juice vs. (B) white grape juice; (A) elderberry juice vs. (B) water; (A) 15% sucrose vs. (B) elderberry juice.

Training in this task typically took 2.5-4 months and proceeded as described by Kuwabara, Kang, Holy and Padoa-Schioppa ^2^. Briefly, in the first phase of training, animals were presented only one odor in each trial– either octanal (associated with juice A) or octanol (associated with juice B) – on the left or on the right. To obtain the juice, they had to lick the corresponding spout (forced choices). Over sessions, we introduced different odor concentrations corresponding to different juice quantities. Once performance reached 80% correct, we moved to the second phase of training, in which we delivered both odors simultaneously and the animal had to choose between the two options (binary choice). Each training phase typically took 1-1.5 months. Throughout the training period, the animal weight was maintained at 75%-85% of baseline ^39^. Neural recordings started from the beginning of the second phase.

### Logistic analysis of choice data

All the analyses of behavioral and neuronal data were performed in Matlab (MathWorks). Choice patterns were analyzed using logistic regressions ^32^. For each session, we focused on binary choice trials (i.e., we excluded forced choices) and we built the following logistic model:

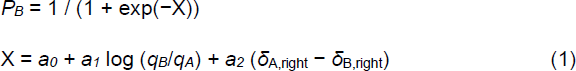

In **Eq.1**, *P_B_* is the probability of choosing juice B; *q_A_* and *q_B_* are the offered quantities; *δ*_A,right_ = 1 if juice A was offered on the right and 0 otherwise, and *δ*_B,right_ = 1 – *δ*_A,right_. From the fitted parameters *a_0_* and *a_1_*, we computed the relative value of the two juices *ρ* = exp(−*a_0_/a_1_*), the choice accuracy *η* = *a_1_*, and the side bias *ε* = – *a_2_*/*a*_1_. Intuitively, *ρ* is the quantity ratio *q_B_*/*q_A_* that makes the animal indifferent between the two juices (**Fig.1C**). The parameter *η* captures the choice accuracy (i.e., consistency); it is proportional to the sigmoid steepness and inversely related to choice variability. The parameter *ε* quantifies a bias favoring one of the two sides; specifically, *ε*<0 (*ε*>0) indicates a choice bias favoring the offer presented on the left (right).

In general, the side bias measured in our experiments was relatively modest (|*ε*| < *ρ*). Thus in the analysis of neuronal activity, we defined candidate variables based on the relative value, disregarding the side bias. To increase our statistical power in computing these variables, we built a reduced logistic model:

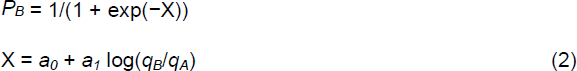

From parameters fitted with **Eq.2**, we computed the relative value *ρ* = exp(−*a_0_/a_1_*) and the sigmoid steepness *η* = *a_1_*. However, for some of the analyses, we imposed restrictive criteria, including only sessions with |*ε*| ≤ 1 (cosine similarity analysis and odd ratios analysis; **Fig.5** and **Fig.7**) or |*ε*| ≤ 0.5 (population analysis; **Fig.7**).

### Longitudinal analysis of choice behavior

We conducted a longitudinal analysis of choice behavior. For each mouse, we counted days starting with the first in which the animal chose between two different juices (T = 0). For each session, we derived the parameters *ρ*, *η*, and *ε* from the logistic fit (**Eq.1**). For each animal and each parameter, we removed outliers (i.e., data points that differed >3 standard deviations from the mean, see **Fig.2D**), and we fitted the remaining values with an exponential function:

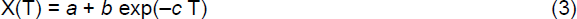

In **Eq.3**, T indicates the day of the session and X = *ρ*, *η*, or *ε*. For each fitted parameter *a*, *b*, and *c*, we obtained a 95% confidence interval. Fitted parameter *a* captured the plateau value (for T→∞, X→*a*). If the 95% confidence interval for either parameter *b* or *c* did not include 0, there was evidence for a significant longitudinal trend; in this case, *c* was the inverse time constant of this trend.

We performed this analysis for each animal and each behavioral parameter *ρ*, *η*, and *ε*. One animal (M1062) was excluded from the analysis because we only had data from 4 sessions. We also conducted a population analysis focusing on the choice accuracy *η* and pooling sessions from all 13 mice. We removed outliers (data points that differed >3 standard deviations from the mean) and we fitted the whole data set with a single exponential function (**Eq.3**).

### Procedures for two-photon calcium imaging

2P Ca^2+^ imaging was performed with an Ultima system (Prairie Technologies) built on an Olympus BX61W1 microscope. A mode-locked laser (Mai Tai DeepSee Ti:Sapphire laser, Spectra-Physics) tuned to 920 nm was raster scanned at 7-11 Hz for excitation while emitted GCaMP6f fluorescence was collected through a green filter (525/70 nm). Laser power at the sample was 20-60 mW. Dwell time was set to 3.2-4 μs. To increase the imaging speed, resolution along the *y*-axis was reduced by a factor of 2. The final pixel size was 2.29x4.58 μm. We used a 20 × 0.45 NA objective (Olympus LCPLN20XIR objective). Thus in each animal we could image a cylinder of 850 μm diameter and 350 μm depth. By varying the focal depth, we could image from multiple fields of view (FOVs). For data collection, we restricted the imaging to depths between 120-370 μm, with 90% of FOVs placed between 180-330 μm. Within a FOV, we often selected a smaller region of interest, to achieve an imaging frequency of ≥7 Hz.

Throughout this paper, we refer to this region of interest as the FOV.

### Cell segmentation and session registration

Cell segmentation was conducted using the CaImAn package ^25^. First, we performed a frame-to-frame non-rigid motion correction to remove potential movement of individual neural signals. Then we extracted and estimated the neuronal fluorescent signal with constrained non-negative matrix factorization (CNMF). We set the criteria to identify a neuron: the signal-to-noise ratio should be >1. After that, we manually checked individual neurons and excluded those without a donut-shaped cell morphology. This manual checking process is blind of cell’s functional property, but only relied on the shape of its spatial footprint. For each cell identified with these procedures, we normalized the fluorescence trace F at each time point to a fixed baseline generated from the sum of the 20^th^ percentile of the average F signal and background signal (generated from CNMF) within each cell region and within a 200 frames time window. This normalization resulted in the ΔF/F signal, which was used for all subsequent analyses.

Data for this study came from 13 mice and 77 FOVs. Each FOV was recorded in 2-16 sessions (mean = 5.95, std = 2.64). For each FOV and each pair of sessions, we registered images and matched neurons using a Bayesian procedure ^26^. First, we aligned the images of the two FOVs by projecting their neuronal footprints onto the same image and finding the rotations and translations that yield the highest cross-correlation between their projections. Then we modeled the distributions of centroid distances and spatial correlations as a weighted sum of the distributions of two subpopulations of cell-pairs, representing same cells and different cells.

Using Bayes’ rule, we obtained the probability (P_same_) for any pair of neighboring cells from different sessions to be the same cell, given their spatial correlation and centroid distance. Finally, we imposed a threshold of P_same_>0.5. Cell pairs that satisfied this criterion were identified as matching. Using these procedures, we typically found that 60%-80% of cells from any given session could be matched with cells recorded in the same FOV in another session.

### Cosine similarity analysis of activity profiles

Referring to the choice task, an offer type was defined by two offered quantities (*q_A_*, *q_B_*); a trial type was defined by two offered quantities, their spatial configuration, and a choice. Because neural activity was recorded at different imaging frequency in different sessions, for the analysis of activity profiles, we first interpolated each cell’s time course and resampled it at 20 Hz. We focused on the ΔF/F signal recorded in two time windows aligned with the offer onset (–600 ms, +1600 ms) and the first lick (–600 ms, +600 ms). For each trial, we joined these two time windows and obtained a single trace. We then averaged traces across trials for each trial type. The activity profile was defined as the concatenation of mean traces obtained for different trial types. Importantly, when comparing the activity profiles recorded for one cell in two sessions, we only included trial types present in both sessions.

The likeness of activity profiles was quantified using cosine similarity (CS). For any neuron *i* recorded in sessions 1 and 2, we indicated the two activity profiles as *AP_i,1_* and *AP_i,2_*, respectively. The CS was defined as:

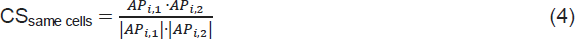

where “·” indicates a vector product and “|x|” indicates the Euclidean norm of the vector x.

Because our measures of ΔF/F were mostly >0, AP vectors were effectively confined to a single orthant of a high-dimensional space. Consequently, CS was expected to be high. To obtain chance level estimates for CS, we generated two benchmark treatments. First, given two sessions 1 and 2, we randomly paired each neuron of session 1 with a neuron of session 2 (treatment: mismatched neurons). Indicating with *i* and *j* two randomly paired neurons in the FOV, we computed:

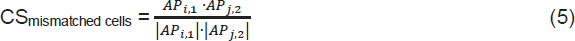

We repeated this operation 10,000 times and we obtained a distribution for CS_mismatched cells_. Second, for each (matched) neuron *i*, we shuffled the activity profile in session 2 by randomly permuting the vector components (treatment: shuffled times). We then computed:

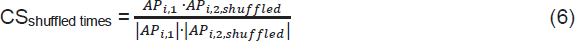

We repeated this operation 10,000 times for each cell, and we obtained a distribution for CS_shuffled times_.

For longitudinal analyses, we counted days for each animal starting the first day in which they chose between different juices (see above). Recordings continued for 12-41 weeks, depending on the animal. For any two sessions 1 and 2, we indicate with T1 and T2 the days in which they took place (always T1 ≤ T2). We also defined ΔT = T2 – T1. On some occasions, we ran two sessions back-to-back in the same day (ΔT = 0). For a longitudinal analysis of activity profiles, we examined CS as a function of the experience in the task (T1) and the time difference between two sessions (ΔT). We partitioned the (T1, ΔT) parameter space in 2-weeks x 2-weeks bins. For given T1 and ΔT, we considered the distribution of CSs obtained across animals and FOVs, and we computed the median(CS). Thus we obtained values of median(CS) as a function of T1 and ΔT. These operations were repeated for the three treatments (same neurons, mismatched neurons, shuffled times). Statistical comparisons between distributions of CSs were conducted using the Kruskal-Wallis test. To increase the statistical power, for some analyses we defined larger time intervals (6-weeks x 6-weeks), referred to as Bins (see **Results**).

### Neuronal encoding of decision variables

We conducted a series of analyses to identify the variables encoded by individual neurons in OFC using essentially the same procedures previously adopted for the analysis of spike counts ^2, 7^. For these analyses, we selected one session per FOV based on two criteria: (1) we only considered sessions with |*ε* | ≤ 0.5 and (2) among these sessions, we selected the one with the largest number of recorded cells. Neural activity (ΔF/F) was examined in five time windows: pre-offer (0.6–0 s before the offer onset), post-offer (0.4–1 s after the offer onset), late delay (1–1.6 s after the offer onset), pre-juice (0.6–0 s before juice delivery onset), post-juice (0–0.6 s after juice delivery onset) (**Fig.1B**). Again, a trial type was defined by two offered quantities, their spatial configuration, and a choice. For each cell, each time window, and each trial type, we averaged the ΔF/F activity the time window and over trials. A neuronal response was defined as the average activity of one neuron in one time window as a function of the trial type.

Task-related responses were identified with an ANOVA (factor: trial type; p<0.01). Neurons that passed this criterion in at least one time window were identified as task-related and included in subsequent analyses. Following previous studies, we defined twelve candidate variables that neurons in OFC might encode (**Supplemental Table 2**). We performed a linear regression of each response passing the ANOVA criterion on each variable, and obtained the corresponding R^2^. If the regression slope differed significantly from zero (p<0.05), the variable was said to explain the response. If the regression slope was indistinguishable from zero, we arbitrarily set the corresponding R^2^ to zero. Neuronal responses were often explained by more than one variable; the variable providing the best fit was that with the highest R^2^.

To identify a limited number of variables that could explain most of the neuronal population, we proceeded with a variable selection analysis. As in previous studies, we used a stepwise procedure and a best-subset procedure ^7, 27^. The stepwise procedure selects at each iteration the variable providing the maximum explanatory power, considering time windows separately. Once a variable is selected, all the responses explained by that variable (including those that would be explained by some other variable with a higher R^2^) are removed from the data set. The residual data set is then examined, and a new variable is selected. The marginal explanatory power of a variable is defined as the fraction of responses that are explained by that variable and that are not explained by any other selected variable. The algorithm continues until the marginal explanatory power of newly selected variables is <2% of the total. Importantly, at each iteration we verify that the marginal explanatory power of each selected variable satisfies this criterion. Previously selected variables whose marginal explanatory power drops below 2% in the presence of a newly selected variable would be eliminated from the selected subset. In practice, however, this situation did not occur for the present data.

Importantly, the stepwise procedure does not guarantee optimality. In contrast, the best-subset procedure is exhaustive. For k = 2, 3, …, the procedure examines all the possible subsets of k variables, computes the total explanatory power (i.e., the number of responses collectively explained), and then selects the subset providing the maximum explanatory power. In this study, the two procedures yielded the same results.

### Stability in the representation of decision variables: Analysis of odds ratios

We conducted a series of analyses to assess whether the variable encoded by any given cell remained stable over extended periods of time. Intuitively, our approach was as follows. For each FOV and any two sessions, we focused on cells identified (matching) and task-related in both sessions. For each cell, we considered variables encoded in the two sessions; if in one session the cell was task-related but did not encode any variable, the cell was classified as *untuned* for that session. On this basis, we constructed for each FOV and for any two sessions a contingency table where rows and columns corresponded to 8 possible classes (7 variables plus *untuned*) recorded in the two sessions, and entries indicated cell counts. In this table, neurons with consistent encoding populated the main diagonal.

Based on this general idea, we conducted a population analysis as follows. First, for each FOV in our data set, we selected two sessions based on two criteria: (1) we imposed |*ε*| ≤ 1, and (2) we selected the two sessions with the largest number of matching, task-related cells. Second, we pooled data across FOVs and examined the resulting table of cell counts (**Fig.7A**). Third, to assess whether cell counts measured for any pair of variables was above or below chance level, we computed the corresponding table of odds ratios (OR) ^40^. Indicating with *C* the table of cell counts, for any entry (*p*, *q*), OR was defined as follows:

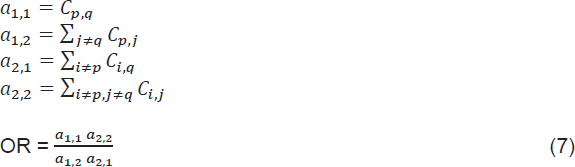

We computed OR for each entry and obtained the full table (**Fig.7B**). Importantly, for each entry, the chance level was OR = 1. Conversely, OR > 1 (OR < 1) indicated that the cell count for that entry was above (below) chance. For each entry, we used Fisher’s exact test (two tails) ^41^ to assess whether the departure from chance was statistically significant.

### Analysis of diagonalization index

The degree of stability in the neuronal representation of decision variables was captured by the relative weight of the diagonal in the OR table. To quantify this weight, we used the diagonalization index (DI), defined as follows. For a *d* x *d* nonnegative matrix *A*,

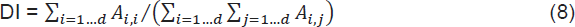

Of note, DI ranges between 0 and 1, and DI = 1 corresponds to a perfectly diagonal matrix – i.e., faultless stability. More generally, the closer DI is to 1, the more stable is the representation.

If all variables were encoded by a similar fraction of cells, the chance level for DI would be 1/n, where n is the number of variables (including *untuned*). However, different variables were encoded with different frequencies. Thus to obtain chance level estimates for DI, we generated a control measurement. Given two sessions 1 and 2, we randomly paired each neuron of session 1 with a neuron of session 2 (treatment: mismatched cells). We then generated the contingency table and calculated DI. We repeated this procedure 10,000 times and we averaged the DIs. For longitudinal analyses, we repeated this procedure for each pair of sessions.

---

**Supplemental Table 1.** Longitudinal analysis of behavior, by animal. For each animal, each behavioral parameter (*ρ*, *ƞ* and *ε*), and each parameter of the exponential fit (*a*, *b*, *c*, see **Eq.3**), the table lists the fitted value and the width of the 95% confidence interval (uncorrected; in parenthesis). When the 95% confidence interval for either parameter *b* or *c* did not include 0, there was evidence of a significant longitudinal trend. Of the 12 animals in which we could conduct this analysis, 6 showed significant longitudinal trends (M0031, M0032, M0036, M1039, M1050, M1053; indicated with **). For 10 of 12 animals, the choice accuracy increased over time (*b*<0), although the trend did not always reach significance level (indicated with *). In some cases, the exponential fit returned a NaN. For parameters b and c, NaN indicated a lack of temporal trend. For parameter a, NaN indicated that the behavioral parameter converged to 0. M1062 was excluded from this analysis because only 4 sessions were available.

| | Relative value ($\rho$) | | | Choice accuracy ($\eta$) | | | Side bias ($\varepsilon$) | | |
| --- | --- | --- | --- | --- | --- | --- | --- | --- | --- |
| Mouse | *a* | *b* | *c* | *a* | *b* | *c* | *a* | *b* | *c* |
| M0031 | 1.92<br>(0.91) | -1.20**<br>(0.78) | 0.02<br>(0.04) | 2.21<br>(2.04) | -1.55*<br>(1.72) | 0.02<br>(0.05) | 3.33<br>(128.3) | -5.22<br>(127.0) | 4 10<sup>-3</sup><br>(0.11) |
| M0032 | 1.56<br>(0.33) | 2.29**<br>(0.98) | 0.07**<br>(0.06) | 2.21<br>(0.44) | -1.5**<br>(0.96) | 0.05<br>(0.07) | NaN | -1.76**<br>(0.89) | 0.10**<br>(0.08) |
| M0036 | 3 10<sup>-6</sup><br>(416.3) | 2.05<br>(415.8) | 9 10<sup>-4</sup><br>(0.19) | 2.23<br>(0.42) | -1.43**<br>(1.34) | 0.09<br>(0.17) | 3 10<sup>-8</sup><br>(0.19) | NaN | NaN |
| M1027 | 1.78<br>(0.19) | -0.22<br>(0.73) | 0.10<br>(2.4) | 1.61<br>(0.19) | -0.66*<br>(0.70) | 0.10<br>(0.75) | 0.09<br>(0.18) | -0.11<br>(0.87) | 20.3<br>(5 10<sup>9</sup>) |
| M1035 | 1.76<br>(0.75) | -1.11<br>(1.81) | 0.08<br>(0.29) | 1.54<br>(0.87) | -0.29*<br>(0.85) | 0.02<br>(0.28) | 28.6<br>(3 10<sup>5</sup>) | -28.4<br>(3 10<sup>5</sup>) | 4 10<sup>-5</sup><br>(0.34) |
| M1039 | 2.82<br>(0.22) | NaN | NaN | 2.51<br>(0.54) | -1.65**<br>(1.02) | 0.01<br>(0.02) | NaN | 0.59<br>(0.62) | 0.02<br>(0.02) |
| M1050 | 1.16<br>(0.14) | -0.32<br>(0.36) | 0.03<br>(0.08) | 2.05<br>(0.23) | -0.92*<br>(1.54) | 0.29<br>(0.88) | NaN | -1.11**<br>(1.01) | 0.18<br>(0.22) |
| M1051 | 1.37<br>(0.13) | -0.12<br>(0.63) | 0.33<br>(7.56) | 1.91<br>(0.31) | NaN | NaN | 0.43<br>(0.18) | NaN | NaN |
| M1052 | 6.08<br>(3.35) | NaN | NaN | 140.2<br>(2 10<sup>5</sup>) | -139.6*<br>(2 10<sup>5</sup>) | 2.64<br>(10<sup>-5</sup>) | 0.56<br>(0.66) | -1.50<br>(3.22) | 0.03<br>(0.10) |
| M1053 | 1.19<br>(0.10) | 0.27<br>(0.45) | 0.06<br>(0.25) | 1.9<br>(0.26) | NaN | NaN | 0.58<br>(0.31) | -0.81**<br>(0.68) | 0.01<br>(0.02) |
| M1057 | 4.07<br>(1.59) | -5 10<sup>6</sup><br>(4 10<sup>9</sup>) | 0.34<br>(17.5) | 1.56<br>(0.42) | -478*<br>(9 10<sup>3</sup>) | 0.14<br>(0.44) | 0.34<br>(0.81) | -110.5<br>(903.8) | 0.09<br>(0.19) |
| M1058 | 1.74<br>(0.95) | -1.13<br>(1.37) | 0.02<br>(0.07) | 1.53<br>(0.29) | -0.19*<br>(1.22) | 0.36<br>(6.36) | 184.0<br>(8 10<sup>5</sup>) | -184.3<br>(8 10<sup>5</sup>) | 1 10<sup>-5</sup><br>( ) |
| M1062 | Excluded from this analysis (only 4 sessions available) |  |  |  |  |  |  |  |  |

**Supplemental Table 2.** Candidate variables. In the analysis of neuronal data, we examined 12 variables. All values were expressed in units of juice B. The relative value *ρ* used to compute the unitary value of juice A was derived in each session from the logistic fit (**Eq.2**).

|  | Candidate variable | Definition |
| --- | --- | --- |
| 1 | Position of A | Left or right position of juice A; 1 if juice A is on left side |
| 2 | Offer value A | Value of juice A offered = $\rho q_A$ |
| 3 | Offer value B | Value of juice B offered = $q_B$ |
| 4 | Offer value ipsi | Value of offer on the ipsilateral side |
| 5 | Offer value contra | Value of offer on the contralateral side |
| 6 | Chosen value | Value of chosen offer |
| 7 | Chosen value A | Chosen value if juice A is chosen, 0 otherwise |
| 8 | Chosen value B | Chosen value if juice B is chosen, 0 otherwise |
| 9 | Chosen value ipsi | Chosen value if ipsilateral offer is chosen, 0 otherwise |
| 10 | Chosen value contra | Chosen value if contralateral offer is chosen, 0 otherwise |
| 11 | Chosen juice | 1 if juice A is chosen; 0 if juice B is chosen |
| 12 | Chosen side | 1 if ipsilateral offer is chosen; 0 if contralateral offer is chosen |

**Supplemental Table 3.** Imaging target locations. Target locations in OFC for each of the 13 mice included in this study. For columns 3-5, distance is in reference to Bregma. The last column details the depth correction introduced to account for the working distance of the GRIN lenses. All lenses were placed in the left hemisphere.

| Mouse ID | Sex | AP (mm) | ML (mm) | DV (mm) | DV correction (mm) |
| --- | --- | --- | --- | --- | --- |
| M0031 | M | 2.6 | -1.55 | -3.3 | 0.15 |
| M0032 | M | 2.6 | -1.55 | -3.3 | 0.15 |
| M0036 | M | 2.6 | -1.55 | -3.1 | 0.15 |
| M1027 | F | 2.6 | -1.3 | -2.6 | 0.25 |
| M1035 | F | 2.7 | -1.4 | -2.6 | 0.3 |
| M1039 | F | 2.7 | -1.35 | -2.5 | 0.3 |
| M1050 | M | 2.8 | -1.3 | -2.6 | 0.3 |
| M1051 | F | 2.7 | -1.4 | -2.65 | 0.3 |
| M1052 | F | 2.8 | -1.5 | -2.65 | 0.3 |
| M1053 | M | 2.6 | -1.6 | -2.65 | 0.3 |
| M1057 | F | 2.7 | -1.6 | -2.65 | 0.3 |
| M1058 | F | 2.7 | -1.6 | -2.65 | 0.3 |
| M1062 | M | 2.7 | -1.7 | -2.6 | 0.25 |

**Supplemental Figure 1.**
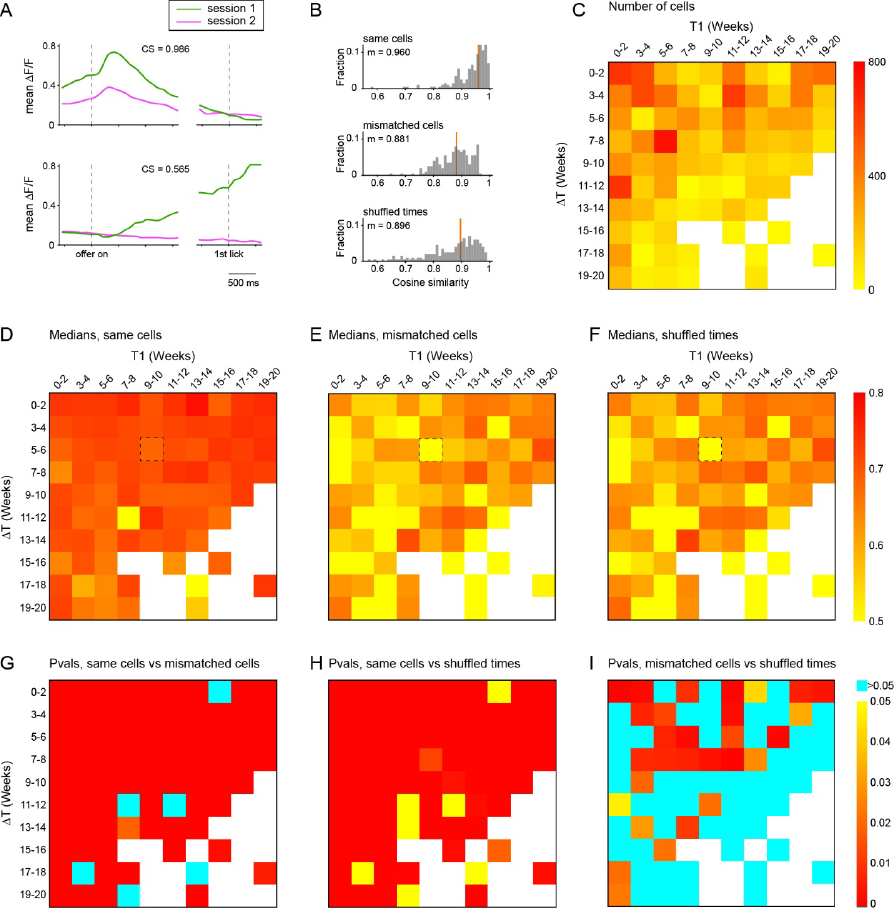
Cosine similarity analysis of activity profiles (alternative procedure). The analyses presented here are similar to those in Fig.4, except that the activity profiles were computed averaging all the trials in one session. **A.** Activity profiles for two example cells. The two neurons were recorded in the same FOV, on day 1 (session 1) and on day 103 (session 2). For each session, we averaged ΔF/F traces across trials in two time windows aligned with offer onset (–600 ms, 1600 ms) and the time of first lick (–600 ms, 600 ms). For each cell, green and pink traces refer to session 1 and session 2, respectively. For cell 1, we measured CS = 0.99. For cell 2, we measured CS = 0.56. **B.** Cosine similarity distributions for the three treatments. Here T1 = weeks 9-10 and ΔT = weeks 5-6. For each distribution, an orange vertical line indicates the median. We measured median(CS_same_) = 0.91, median(CS_mismatched_) = 0.89 and median(CS_shuffled_) = 0.88. **C.** Number of cells matched across sessions as a function of T1 and ΔT. Same as Fig.4E. **DEF.** Measures of median(CS) as a function of T1 and ΔT. All conventions as in Fig. 4FGH. The black dashed rectangle highlights the bin illustrated in panel **B**. **GHI.** Statistical comparison of different treatments. All conventions as in Fig.4IJK. All p-values were obtained from a Kruskal-Wallis test.

**Supplemental Figure 2.**
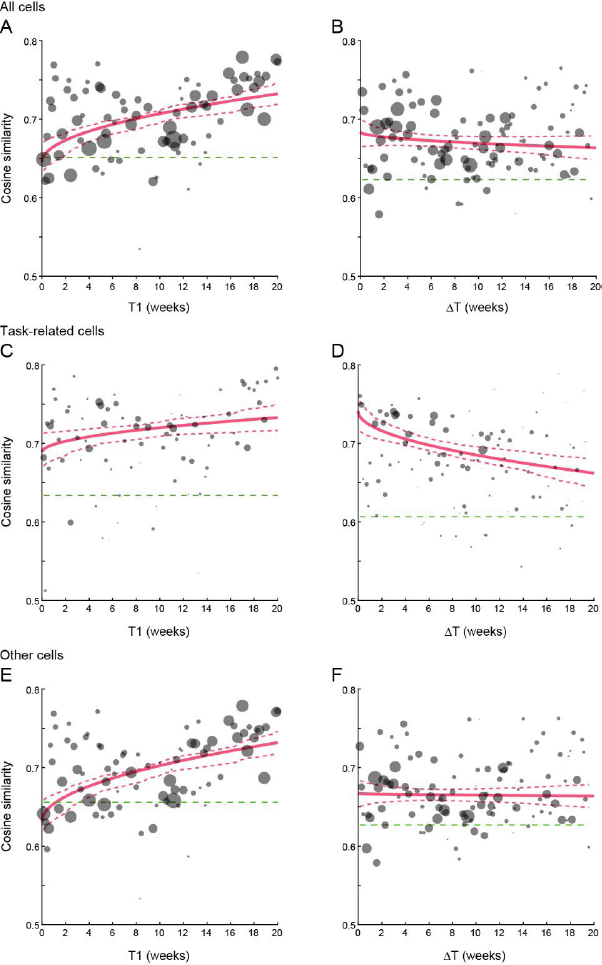
Cosine similarity, longitudinal trends, and task-relatedness. The longitudinal analysis presented in Fig.5 indicated that activity profiles became more reproducible as the animals became more experienced in the task (i.e., as a function of T1; Fig.5B). Conversely, activity profiles for any given neuron changed somewhat as a function of time passage (ΔT; Fig.5C). This phenomenon, termed representational drift, was statistically significant but quantitatively modest. Indeed, the time necessary for a full reorganization was estimated at 3.5 years – i.e., longer than the animal lifetime. One concern was whether the representational drift would be more pronounced specifically in task-related cells. Thus we repeated the analyses of Fig.5 separately for task-related cells and other cells. **AB.** All cells. Same as in Fig.5BC. **CD.** Task-related cells. **EF.** Other cells. Interestingly, CS was higher for task-related cells (panel **C**) than for other cells (panel **E**) starting from T1 = 0. Restricting the representational drift analysis to task-related cells (panel **D**), we fitted the model CS(ΔT) = *a_0_* + *a_1_* ΔT^1/2^, and we obtained the values *a_0_* = 0.7411 and *a_1_* = -6.6693 10^-3^ day^-1/2^. For the same population, we computed *b* = median(CS_mismatched cells_) = 0.606575 (green dashed line). On these bases, we estimated the time necessary for full reorganization ΔT_FR_ = 407 days. Thus, even under the assumption that the reorganization involves exclusively task-related cells, the time to full reorganization would be >1 year – i.e., more than twice the time course of our longest experiments. In other words, for practical purposes, activity profiles can be considered longitudinally stable.

**Supplemental Figure 3.**
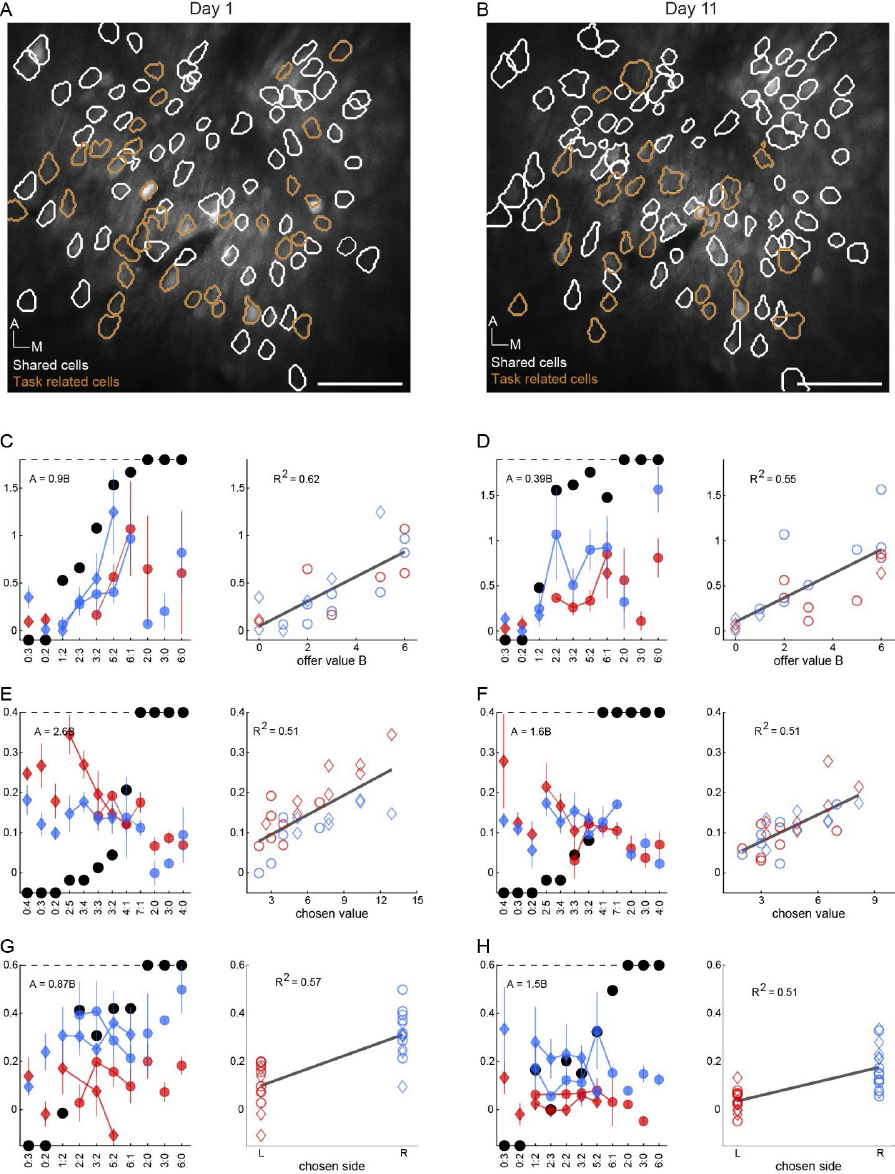
Encoding stability, example cells. **AB.** Example FOVs recorded on day 1 (A) and day 11 (B). Orange contours highlight cells that were both shared and task-related (day 1: N = 24; day 11: N = 33). White contours highlight cells that were shared but not task-related (day 1: N = 74; day 11: N = 65). Anterior and medial directions are indicated; the scale bar indicates 100 μm. **CD.** Example cell the *offer value B* in both sessions. All conventions are as in Fig.6A. **EF.** Example cell encoding the *chosen value* in both sessions. **GH.** Example cell encoding the *chosen side* in both sessions.

**Supplemental Figure 4.**
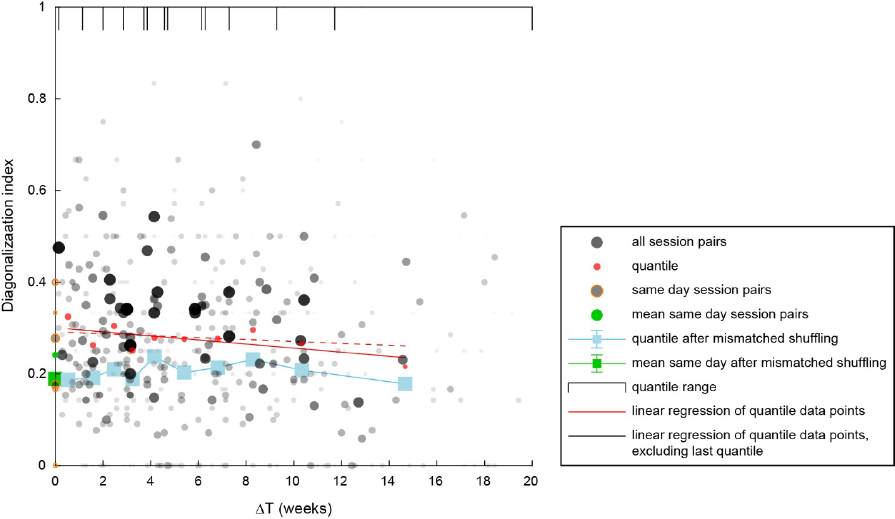
Longitudinal analysis of encoding stability. For each FOV and for each pair of sessions, we generated a contingency table and an OR table similar to those displayed in Fig.7. We quantified the diagonal nature of the OR table using the diagonalization index (DI; **Methods**, **Eq.8**), and we examined how DI varied as a function of ΔT. The figure illustrates the entire data set. The x-and y-axis are ΔT and DI, respectively, and each data point represents one pair of sessions. The gray shade and size of the data point represent the number of neurons contributing to the data point (matched, task-related cells). In general, DI varied substantially from session pair to session pair – across the whole data set, mean(DI) = 0.26 and std(DI) = 0.21. A simple weighted linear regression (with weights based on number of cells) indicated that DI did not significantly decrease as a function of ΔT (p = 0.059, F test). We also divided sessions in 10 quantiles (each with the same number of cells). For each quantile, we computed the mean(DI) (red dots), and we regressed the resulting data points against ΔT. By this measure, DI significantly decreased as a function of ΔT (p = 0.043, F test). However, this result was driven by the last data point – when the regression was repeated excluding the last data point, the relation was not statistically significant (p = 0.44, F test). Since the longitudinal values of DI seemed relatively low, we examined two benchmarks. First, we randomly (mis)matched neurons of session 1 with neurons of session 2. We repeated this procedure 10,000 times for each session pairs, divided ΔT in 10 quantiles, and then computed the mean(DI_mismatched cells_) for each quantile (blue squares). Comparing the quantile measures obtained for actual data (same neurons) and for mismatched neurons we found the mean(DI) for actual data was significantly higher (p = 5 10^-5^, paired t-test). In other words, for any ΔT, the consistency in classification was significantly higher than that expected by chance. Second, we computed DI for all the available session pairs with ΔT = 0 (repeated sessions in the same day; 6 session pairs, N = 59 cells; orange circles with gray filling). For this data set, we found mean(DI_ΔT=0_) = 0.19 (green dot), std(DI_ΔT=0_) = 0.15. The fact that this measure was close to that obtained for the whole data set supported the notion that the encoding of decision variables remained very stable over time. Concurrently, it suggested that our classification procedures were noisy (see main text).

## Acknowledgements

We thank members of the Padoa-Schioppa lab and the Holy Lab for helpful discussions. This work was supported by the National Institutes of Health (grants number R01-DA055709, R01-DA032758 and R21-DA042882 to CPS and R01-DC020034 to TEH).

## Contributions

M.Z., A.L., T.E.H. and C.P.S. designed the study. M.Z. and A.L. performed the experiments and collected the data with assistance from M.C. M.Z. and A.L. processed and analyzed the data. H.S. and M.C. helped with animal training. A.B. helped with anatomy and histology. C.P.S and T.E.H. supervised the project. M.Z. wrote the initial draft. All authors edited the manuscript.

